# Mutagenic mechanisms of cancer-associated DNA polymerase ε alleles

**DOI:** 10.1101/2020.09.04.270124

**Authors:** Mareike Herzog, Elisa Alonso-Perez, Israel Salguero, Jonas Warringer, David J. Adams, Stephen P. Jackson, Fabio Puddu

**Affiliations:** The Wellcome/Cancer Research UK Gurdon Institute and Department of Biochemistry, University of Cambridge, Tennis Court Road, Cambridge CB2 1QN, UK; The Wellcome Sanger Institute, Hinxton CB10 1HH, UK; Department of Chemistry and Molecular Biology, University of Gothenburg, Medicinaregatan 9 C, 413 90, Göteborg, Sweden

## Abstract

A single amino acid residue change in the exonuclease domain of human DNA polymerase ε, P286R, is associated with the development of colorectal cancers, and has been shown to impart a mutagenic phenotype. Perhaps unexpectedly, the corresponding Pol ε allele in the yeast *Saccharomyces cerevisiae* (*pol2-P301R*), was found to drive greater mutagenesis than exonuclease-deficient Pol ε (*pol2-4*), a phenotype sometimes termed *ultra*-mutagenesis. By studying the impact on mutation frequency, type, replication-strand bias, and sequence context, we show that *ultra*-mutagenesis is commonly observed in cells carrying a range of cancer-associated Pol ε exonuclease domain alleles. Similarities between mutations generated by these alleles and those generated in *pol2-4* cells indicate a shared mechanism of mutagenesis that yields a mutation pattern similar to cancer Signature 14. Comparison of *POL2 ultra*-mutator with *pol2-M644G*, a mutant in the polymerase domain decreasing Pol ε fidelity, revealed unexpected analogies in the sequence context and strand bias of mutations. Analysis of mutational patterns unique to exonuclease domain mutant cells suggests that backtracking of the polymerase, when the mismatched primer end cannot be accommodated in the proofreading domain, results in the observed increase in insertions and T>A mutations in specific sequence contexts.

## INTRODUCTION

Large genomic rearrangements are a common feature of many types of cancer, but widespread hypermutation — the extensive accumulation of single nucleotide variants (SNVs) or small insertion/deletions (INDELs) — is relatively rare[1]. Hypermutation usually arises from exposure to mutagens, such as ultra-violet light or tobacco smoke, and/or from hereditary or acquired DNA repair defects, leaving behind specific mutation signatures [2]. DNA mismatch repair (MMR) inactivation, for example, has long been known to drive somatic hypermutation that leads to a class of hereditary colorectal cancers [3]. More recently, specific mutator alleles in the exonuclease (proofreading) domain of replicative DNA polymerases δ and ε (Pol δ and Pol ε), such as *POLE*-*P286R*, were found to foster SNV hypermutation in the presence of functional MMR, and drive the development of hereditary colorectal or endometrial cancers [4–6].

The proofreading activity of B-family DNA polymerases (such as Pol δ and Pol ε) is triggered by the presence of a base-pair mismatch between the template and the nascent DNA strand at the primer-template junction [7]. In these situations, the 3’ end of the primer is melted and moved to the spatially separate proofreading domain, where one or more nucleotides are exonucleolytically degraded [8–10]. The primer end is then returned to the polymerase domain, where DNA synthesis can continue [11,12].

The origin of hypermutation in cancer cells with Pol ε proofreading domain mutants was originally ascribed to inactivation of Pol ε exonuclease activity [4]. Subsequent work in the yeast *S. cerevisiae*, however, revealed that *pol2-P301R* — the ortholog of the human Pol ε mutation *POLE*-*P286R* — drives substantially greater mutagenesis than Pol ε exo^−^ [13], an exonuclease-deficient variant of Pol ε encoded by the yeast *pol2-4* allele [14]. This indicates that inactivation of the exonuclease activity is not primarily the origin of the massive mutation accumulation observed in *pol2-P301R* cells; and accordingly, biochemical work revealed that exonuclease activity is still detectable in the P301R mutant polymerase (~20% to ~60% of wild-type, depending on the assay) [15]. Further structural analyses revealed that this amino acid residue change creates a barrier at the entrance of the exonuclease domain, possibly preventing the newly synthesized strand from accessing it [16]. Inability to position the mismatched primer in the proofreading domain, and the observed increased mismatch extension ability, could explain the *ultra*-mutagenic phenotype, but *in vitro* polymerase assays failed to recapitulate *ultra*-mutagenesis [15]. These observations opened up the possibility that other cellular processes may be involved in the generation of mutations and motivated the studies we describe herein.

## RESULTS

### A spectrum of DNA Pol ε *ultra*-mutator alleles

As an approach to investigate how *ultra*-mutator Pol ε mutants exert their genotoxic activities *in vivo*, we focused on *POLE* alleles (**Fig. 1A**) originally described as drivers of colorectal and endometrial cancers [4–6]. To avoid confounding factors that would arise from conducting such studies directly in cancer cell lines, such as a higher background of genomic instability, we introduced, where possible, the corresponding mutations — hereafter collectively referred as *pol2-c* — in heterozygous state into the diploid yeast strain W303. Thus, we used site-directed mutagenesis to generate eight *pol2-c* alleles for most of these cancer-associated residues evolutionary conserved between human and yeast Pol ε (**Fig. 1A**). Several colonies (18–54) for each POL2/*pol2-c* heterozygous diploid strain were then independently cultured through single-cell bottlenecks (26 passages; ~500 cell generations), allowing mutations to accumulate in the genome. Mutational events that occurred during the experiment were identified by whole-genome sequencing of each mutation accumulation (MA) line at the start and the end of the experiment. For comparison, we also carried out such analyses of MA lines containing wild-type *POL2* or a proofreading defective *pol2-4* allele [14,17]. Since at each passage colonies were randomly selected, we expect mutation rates and spectra to be unbiased with regard to their consequences on gene function, except for overlooking lethal mutations.

**Figure 1.**
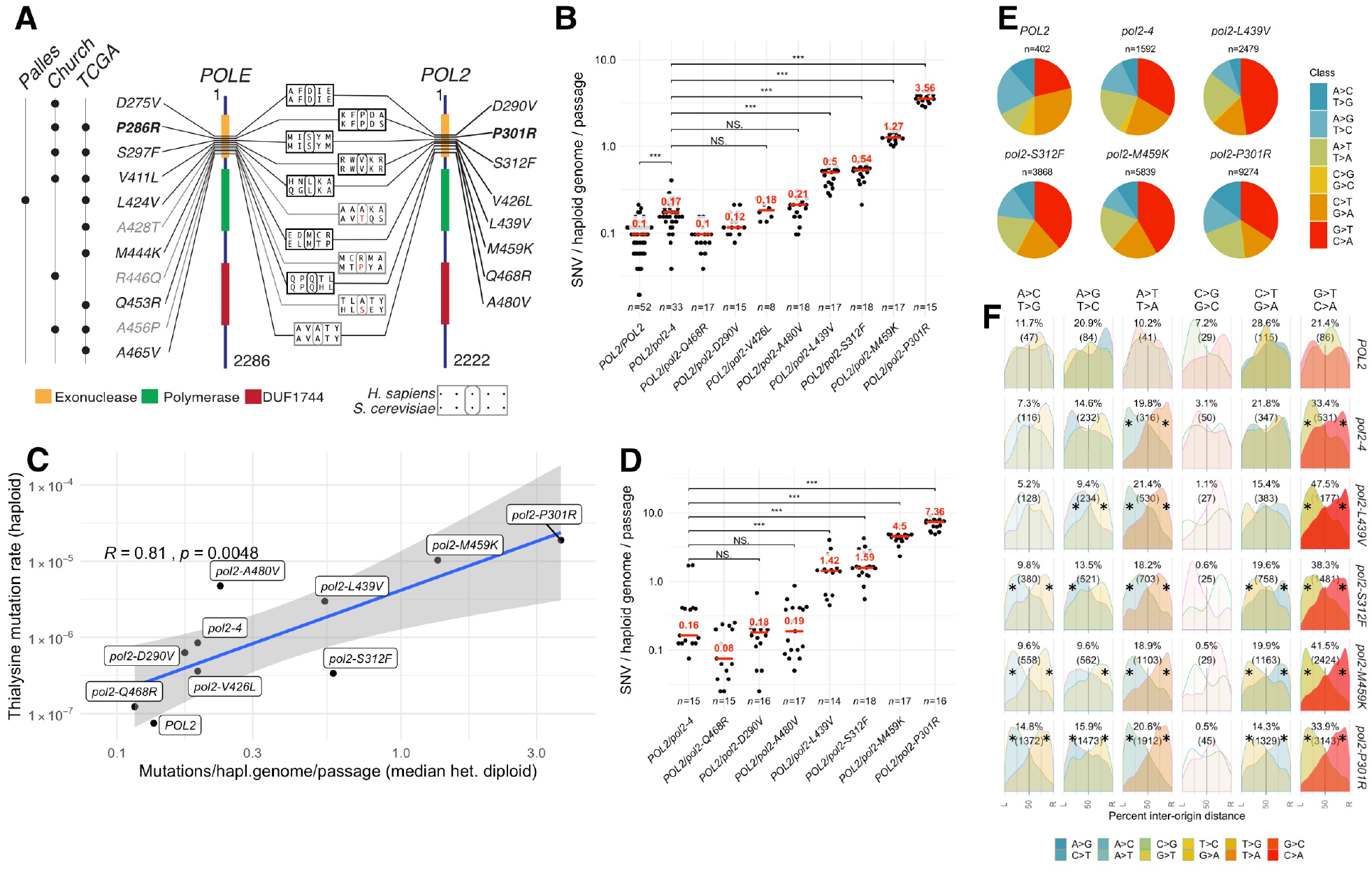
Analysis of *ultra-mutator* alleles of DNA polymerase ε modelled in *S. cerevisiae.* **A.** Outline of DNA polymerase ε ultra-mutator alleles identified in cancers and corresponding mutations in S. *cerevisiae POL2;* the numbers indicate the residue position and domains are indicated with coloured boxes (DUF: domain with unknown function; Palles: mutation first reported in Palles C., *et. al,* Nat. Gen. 2013; Church: mutation first reported in Church DN, *et al.,* Hum. Mol. Genet. 2013; TCGA: mutation present in The Cancer Genome Atlas). Mutations in grey occur in residues not evolutionary conserved in S. *cerevisiae.* The most studied mutation (P286R/P301R) is indicated in bold. **B.** Rates of mutation accumulation in diploid yeast strains carrying indicated heterozygous *POL2* alleles propagated through single-cell passage bottlenecks for ~500 generations. Each independent evolution line is indicated by a dot, while the median is indicated in red. The number of independent lines *(n)* is indicated below. Statistical test: Mann-Whitney-Wilcoxon **C.** Comparison of mutation accumulation rates (x-axis) and loss-of-function mutation rates at the *LYP1* locus (y-axis). Blue line indicates the linear regression model; shaded area the 95% confidence interval; *R* is Pearson’s correlation coefficient. **D.** Rates of mutation accumulation in strains carrying the indicated heterozygous *POL2* alleles propagated through small population passage bottlenecks for 350~450 generations. Different colonies derived from the final population are indicated by dots, while the median is indicated in red. The number of colonies studied *(n)* is indicated below. Statistical test: Mann-Whitney-Wilcoxon. E. Proportion of each mutation type accumulated in the presence of the indicated *pol2* allele. F. Geographical distribution of the density of mutations by type in relation to replication origins. The origins to the left and to the right of each mutation are indicated by L and R respectively, and the distance is expressed as a percentage of the inter-origin distance of each origin pair (L→50%: leading strand; 50%→R: lagging strand). The intensity of the colour is proportional to the frequency of each mutation channel. Asterisks indicate mutation types significantly deviating from uniformity (p<0.01 χ^2^ test)

In line with previous estimates (1.67–3.8 **×** 10^−10^ SNV/generation/bp) [18–20], wild-type MA lines acquired a median of ~0.12 SNV/haploid genome/passage (or ~4.97 **×** 10^−10^ SNV/generation/bp), while strains carrying a proofreading-defective allele (*pol2-4/POL2*) displayed a modest increase over this rate (~30% or 6.6 **×** 10^−10^ SNV/generation/bp; **Fig. 1B)**. Four *pol2-c* alleles led to either no detectable mutator phenotype (Q468R, D290V) or a hypermutator phenotype similar in magnitude to that of *pol2-4/POL2* cells (V426L, A480V; **Fig. 1B**). In contrast, four other Pol ε alleles (P301R, M459K, S312F, L439V) accumulated considerably more mutations than would be expected by simple lack of exonuclease activity (3–23 times or 2.1–15.5 **×** 10^−9^ SNV/generation/bp), thus reflecting an *ultra*-mutator phenotype. Since the growth rates of these strains did not substantially differ from those of the other *POL2* mutants or from the wild-type strain (**Additional file 1: fig. S1A**), MA could be taken as an accurate reflection of mutation rates. Accordingly, when we characterised haploid strains carrying wild-type or various mutant *POL2* alleles for their rates of loss-of-function mutations at the *LYP1* locus (yielding thialysine resistance), there was good concordance (*R* = 0.81) between these data and results from MA experiments (**Fig. 1C**).

Notably, MA in haploid cells was substantially higher than in heterozygotic diploid cells, implying that the presence of a wild-type polymerase reduces the mutagenic effects of the hypermutator allele (**Additional File 1: fig. S1B**). Our results also showed that the presence of a wild-type *POL2* mitigates the effect of *pol2-c* mutants by more than half, suggesting that in diploid cells, the wild-type polymerase is preferentially expressed or used, or that wild-type Pol ε can correct some errors introduced by Pol ε-*ultra* mutants **(Additional File 1: fig. S1B)**. The haploid state also unmasked an *ultra*-mutator phenotype for *pol2-A480V*, which in heterozygosis did not accumulate significantly more mutations than the corresponding exonuclease deficient strain (compare blue and red *p*-values, **Additional File 1: fig. S1B**). In contrast to SNVs, the strongest *pol2-ultra* alleles only led to a small but statistically significant increase in the accumulation of INDELs in haploid cells (**Additional File 1: fig. S1C)**.

Assessing MA in cells propagated through population bottlenecks of ~3 × 10^4^ cells broadly confirmed our initial observations, despite a larger number of mutations per passage, and much higher variability between different colonies (**Fig. 1D**). These effects likely arose from the experimental settings: population expansion through ~10^4^ rather than single cell bottlenecks presumably allowed different clones to grow at different rates, and random sampling at each passage would favour the propagation and analysis of faster-growing clones, which would have completed more DNA replications and therefore accumulated more neutral or adaptive mutations than slower growing ones. The appearance and selection of anti-mutator suppressor mutations [21] could also explain the increased variability in mutation numbers.

Taken together, our results showed that *ultra-*mutagenesis — the accumulation of considerably more mutations than would be expected by loss of exonuclease activity — is a common outcome for mutations in the proofreading domain of Pol ε that are found in cancers.

### Exonuclease deficient and *pol2-ultra* alleles have similar mutational spectra

Analysis of the mutational spectra generated in the absence of Pol ε exonuclease activity (*pol2-4*) revealed a relative increase in the frequency of A>T transversions, which is further expanded in *pol2-ultra* cells, while C>G transversions — the rarest class in wild-type strains — becomes relatively rarer (**Fig. 1E**). Most variants generated in the presence of Pol ε mutants are likely introduced on the replication leading strand, where the activity of this DNA polymerase is confined [22,23]. To measure the replication-strand bias of the observed mutations, we calculated the relative distance of each mutation from the replication origins located to its immediate left and right, and then calculated the mutational density of each complementary mutation pair (e.g. G>A and C>T) as a function of the distance (**Fig. 1F**). This analysis revealed a strong asymmetry (Cohen’s *w*=0.43–0.51) in the distribution of A>C, A>T, and G>T in *pol2-P301R* cells, and a weaker asymmetry (*w*=0.17– 0.29) for A>G and C>T mutations. In particular A>C, A>G, A>T, G>A, and G>T were observed more frequently on the leading strand than their complementary counterparts (**Fig. 1F and Additional File 1: fig. S2A**; the low mutation counts for C>G/G>C transversions did not permit establishment of whether a bias is present in this channel). Strikingly, this pattern of mutagenesis was observed in cells carrying both stronger and weaker ultra-mutator alleles, and in *pol2-4* cells as well, with the degree of asymmetry increasing with the total number of mutations available for analysis (**Additional File 1: fig. S2B**). Taken together, these results strongly suggest a common mechanistic origin for mutations observed in cells lacking Pol ε exonuclease activity and in cells carrying *ultra*-mutator Pol ε variants. They also indicate that SNV accumulation in *pol2-4* cells does not originate from simple lack of exonucleolytic activity.

### Synergism of Pol ε exonuclease domain mutants with MMR deficiency

Mismatch-repair (MMR) recognises different DNA duplex mis-pairs with different efficiencies [24], thereby distorting the frequencies of different mutation classes from the frequencies generated by DNA polymerases. As an approach to determine the mutational patterns as they are generated by Pol ε *ultra*-mutators, we attempted to delete *MSH2*, which encodes a mismatch binding ATPase that is required for all branches of MMR. As previously observed for several other DNA polymerase mutator alleles [15,17,21], sporulation of most *pol2-ultra msh2Δ* heterozygous diploids did not yield viable double mutant strains for the strongest ultra-mutators (*pol2-P301R*, *pol2-M459K*, and *pol2-S312F*). Microscopic observation of these spores revealed that they germinated to vegetative cells but ceased to divide after a few generations, a phenotype consistent with extreme mutational burden leading to “error-induced extinction” [21]. By contrast, double mutants for the weak mutator *pol2-4* in combination with *msh2Δ* were readily obtained from corresponding heterozygous diploids but displayed reduced colony size compared to MMR-proficient controls. We also managed to obtain a viable MMR-deficient version of *pol2*–*L439V—*a relatively weak *ultra*-mutator. MA analyses of these strains confirmed that, similarly to heterozygous diploids, *pol2-4* and *pol2-L439V* accumulated SNVs at a faster rate compared to wild-type strains in the presence of functional MMR (~4X and ~12X respectively; **fig. 2A and Additional File 1: fig. S1B**), while disruption of *MSH2* alone resulted in a ~14-fold increase (or ~7.4**×** 10–9 SNV/bp/generation, similar to a previous estimate of 4.8**×** 10–9 SNV/bp/generation [25]). In contrast, a dramatic increase in SNV accumulation was evident when mismatch repair inactivation was combined with *pol2-4* or *pol2-L439V* (~365X and~840X, respectively; **Fig. 2A**). These numbers are well above the expected mutation rate increases in the double mutants under an additive model (~18X and ~26X respectively), demonstrating a synergistic interaction. Additionally, *pol2-L439V* accumulated more mutations than *pol2-4* even in the absence of MMR, indicating that the ultra-mutator phenotype does not arise from a differential mismatch repair efficiency. Analysis of allele frequencies and status of the mating type (*MAT/HM*) loci also revealed that, unexpectedly, both *msh2Δpol2-4* and *msh2Δpol2-L439V* strains were actually diploid. Both strains were homozygous for *MSH2* deletion and *pol2* mutations, but the former was a *MAT***a**/*MAT***a** diploid— possibly originating from a whole-genome duplication event—and the latter a *MAT***a**/*MAT***alpha** diploid— possibly originating from homothallic mating after a rare mating-type switch event. These results suggest that a transition to the diploid state facilitates survival in the face of extreme mutagenesis as expected from the fact that deleterious mutations are frequently recessive and often masked in a heterozygotic diploid state.

**Figure 2.**
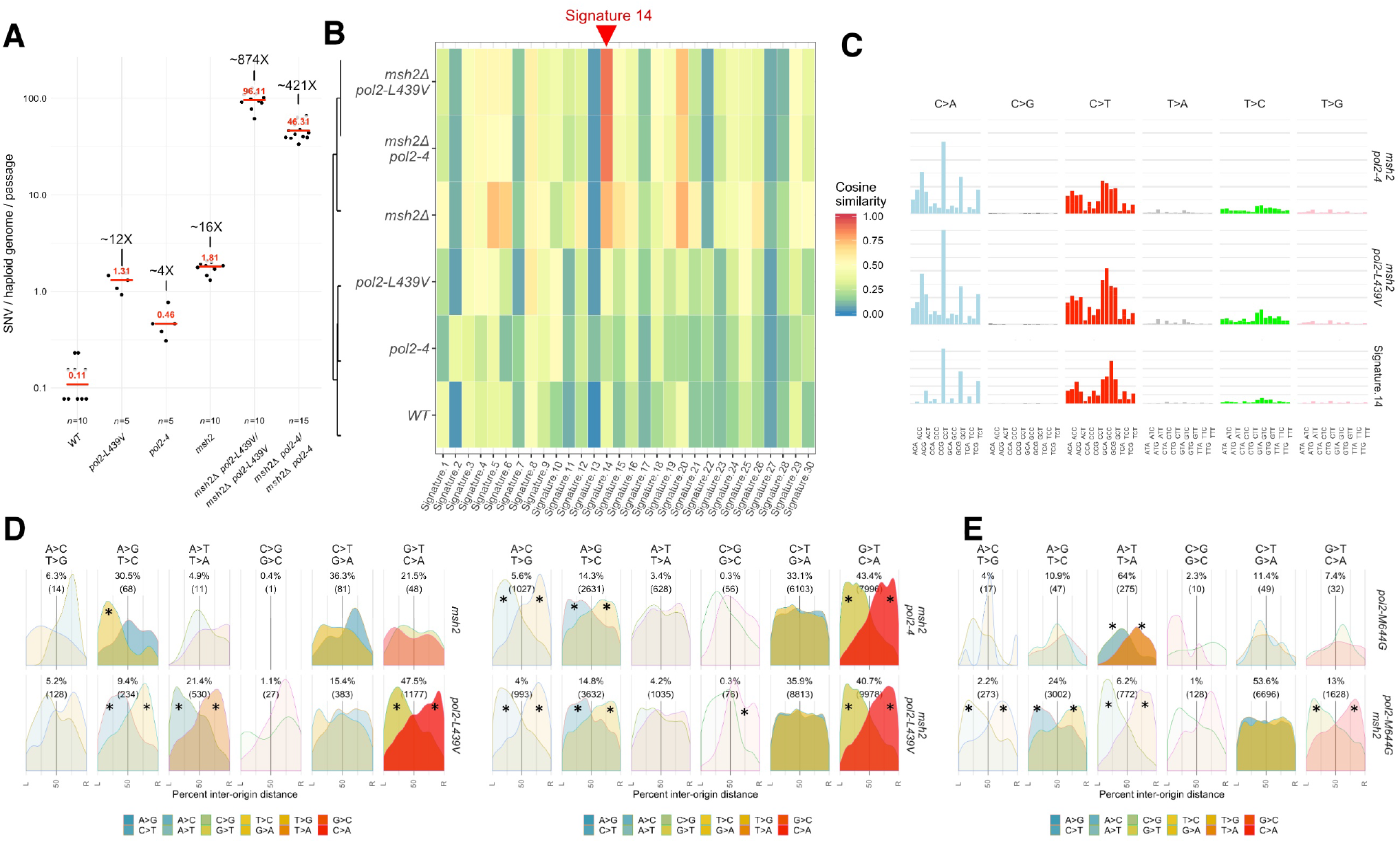
Mutation signatures generated in the presence of Pol ε exonuclease domain alleles. **A.** Number of SNVs generated during single-cell bottleneck propagation by either Pol2 *exo*^−^*(pol2-4)* or the Pol2-ultra mutant L439V in the presence or in the absence of functional mismatch repair. The median is indicated in red. **B.** Similarity between the SNV patterns generated by DNA polymerase mutants in the presence or in the absence of functional mismatch repair and the 30 mutation signatures identified in cancers. Colours indicate cosine similarity (Alexandrov, 2013). **C.** COSMIC-style representation of the mutation patterns generated by Pol2 *exo*^−^*(po/2-4)* or the Pol2-ultra mutant L439V and the most similar COSMIC signature (Signature 14). **D.** Geographical distribution of the density of mutations by type in relation to replication origins. The origins to the left and to the right of each mutation are indicated by L and R, and the distance is expressed as a percentage of the inter-origin distance of each origin pair. The data for *pol2-L439V* have been carried over from Fig. I for comparison purposes. The intensity of each mutation channel is proportional to the intensity of the colour. Asterisks indicate mutation types significantly deviating from uniformity (p<0.01 χ^2^ test). **E.** Geographical distribution of the density of mutations by type in relation to replication origins generated by the *pol2-M644G* allele in the presence or in the absence of functional MMR.

### Pol ε – L439V and Pol ε exo^−^ yield replication-strand biased Signature 14

Analysis of the trinucleotide context in which mutations are introduced by Pol ε – L439V and Pol ε exo^−^ revealed that these share a very similar mutational profile, with striking similarity to COSMIC Signature 14, one of the 30 mutational signatures initially identified in cancers (**Fig. 2B and 2C**). Signature 14 was originally identified in uterine cancers and low grade gliomas [2], and is also observed in cancers carrying both *POLE* mutations and microsatellite instability — the latter a feature of MMR inactivation [26]. Replication-strand bias analysis revealed a strong preference for A>C, A>G, and G>T mutations on the leading strand, as it was observed in the presence of functional MMR (**Fig. 2D and Additional File 1: fig. S2A and S2C**). In this case, the relatively high overall number of mutations we obtained also allowed detection of a preference for G>C mutations on the lagging strand. Conversely, the preference for A>T and G>A mutations on the leading strand was reduced or disappeared when MMR was inactivated (**Fig. 2D and Additional File 1: fig. S2A and S2C**), suggesting that while Pol2 mutants equally introduced these mutations and their complementary ones (e.g. A>T and T>A), MMR corrected one (T>A) more efficiently than the other.

To determine if the observed mutational patterns were specific for Pol ε exonuclease domain mutator alleles, we re-analysed previously published data for the *pol2-M644G* mutant [27], which carries a mutation in the polymerase domain of Pol ε that creates a “looser” active site that allows mis-incorporation of dNTPs and rNTPs [22,28]. In the absence of confounding MMR effects, we found that the activity of each mutation channel was substantially different between exonuclease and polymerase domain mutants. However, the overall replication-strand bias was strikingly similar, with only one major difference in the A>T/T>A channel: while polymerase domain alleles were more likely to produce A>T mutations by mispairing T:dT more frequently than A:dA, exonuclease domain alleles produced both types of mis-pair essentially equally (**Fig. 2E Additional File 1: fig. S2D**). The striking similarity between mutations introduced in the genome by Pol ε exonuclease and polymerase domain mutants suggest that, with some minor differences, a similar mutagenic process is active in cells carrying either mutant.

### A unique signature generated by Pol ε exonuclease domain alleles

We next compared the context in which every class of mutation was observed on the two replication strands. To do this, we pooled all mutations from *pol2-4 msh2Δ* and *pol2-L439V msh2Δ* strains, given their overall similarity (**Additional File 1: fig. S3**) and apparent common origin, and compared them with mutations generated by the polymerase domain allele *pol2-M644G* in the absence of *MSH2* [27]. This indicated that alteration of either the exonuclease or polymerase domain leads to the mis-insertion of dCTP opposite to the second T of a -TT- dimer template; less frequently the inverse is also observed (dTTP mis-insertion opposite to the C of a -TC- dimer template, **Fig. 3A, B red boxes**). Overall, these two classes of mutations were more prevalent in exonuclease-than in polymerase-domain mutator strains (~34% vs. ~10% of all mutations, respectively; p<0.01 χ2 z-test of given proportions) suggesting that a similar mutagenic mechanism occurs with different intensity in different mutants.

**Figure 3.**
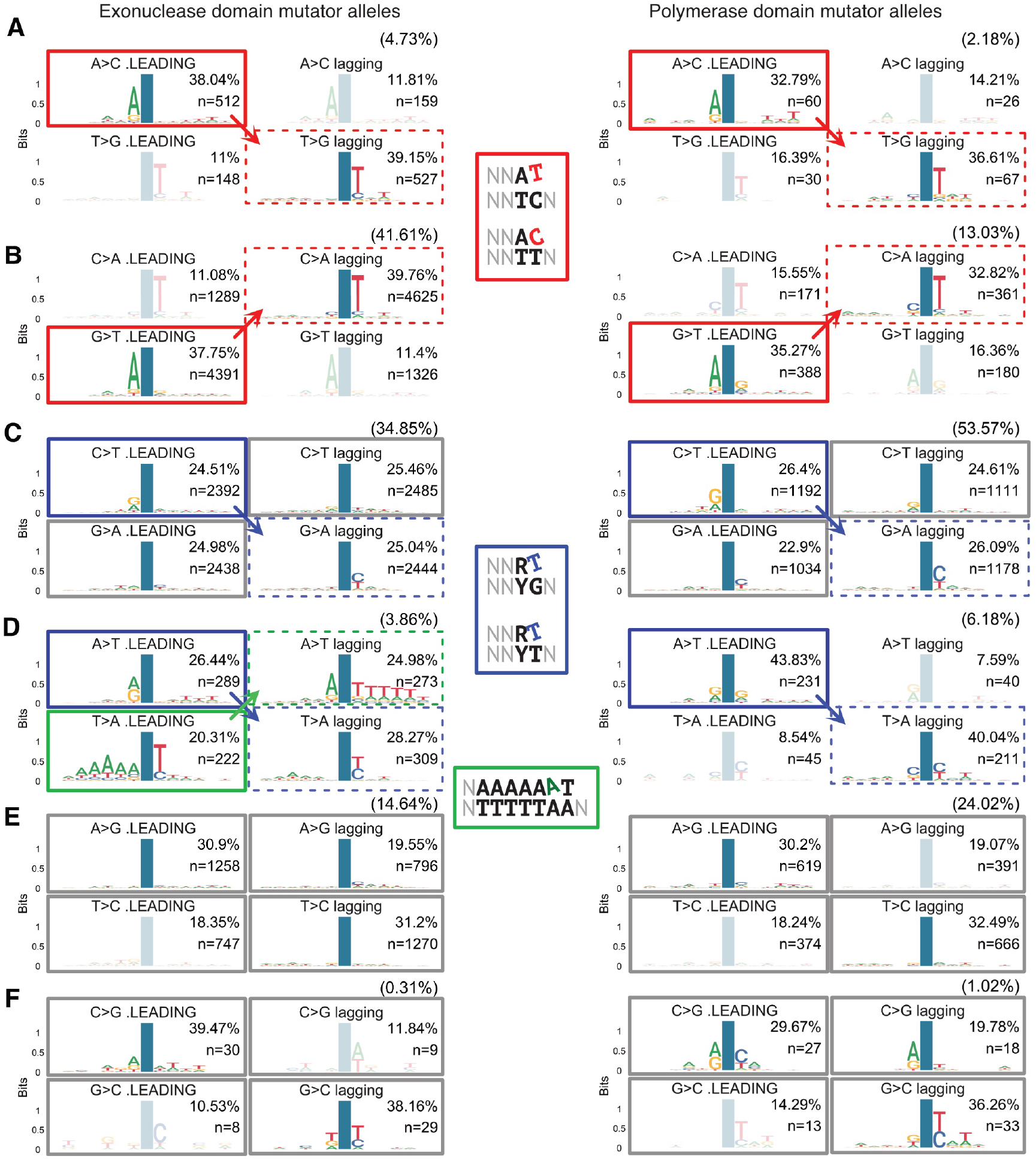
Sequence context of the *de novo* mutations generated in cells with Pol ε exonuclease or polymerase domain mutant alleles. **A-F**. Sequence context in which each *de nova* mutation class occurs in cells carrying exonuclease domain mutator alleles *(pol2-4* or *pol2-L439V)* or polymerase domain mutator alleles *(pol2-M644G),* in the absence of functional MMR *(msh2Δ).* Mutations were categorised by type and by whether they occurred in regions of the Watson strand synthesised as leading or lagging strand. The frequency of each mutation class is indicated in brackets. Within each class the frequency of the four types is indicated in each panel.The position of the mutation is indicated by a blue column. Solid coloured boxes indicate that the mutation observed on the leading strand is the one introduced directly or indirectly by Pol ε on the leading strand. Dashed boxes indicate that the complementary sequence context is observed in lagging strand regions. These arise from Pol ε-mediated synthesis of the Crick strand as leading strand in these regions. Transparent classes indicate mutations that are likely artefacts created by leading strand replication not precisely terminating at the inter-origin midpoint and/or by some origins being passively replicated. Grey boxes indicate mutation classes for which no sequence context can be identified or not enough mutations were studied.

A second shared pattern of mutagenesis between polymerase and exonuclease domain mutants of Pol ε is the mis-insertion of dTTP in front of G or T, which often occurred after a pyrimidine in the template (**Fig. 3C and 3D, blue boxes**). Notably, these classes of mutations were more frequently observed in MA lines with a polymerase-domain mutator allele (~33% of all mutations) than in strains with exonuclease-domain mutator alleles (~19% of all mutations; p<0.01). Other classes of mutation showed very little or no sequence-context specificity, despite their overall relatively high prevalence (**Fig. 3C and 3E; grey boxes**). A notable exception to this was insertions of A in front of the first A after a long T homopolymer that is followed by an AA dimer (TTTTTAA for example; **Fig. 3D, green boxes**). Despite being a relatively uncommon event (~2% of all mutations), this signature was unique to exonuclease-domain mutators, being completely absent from the mutational spectra of *pol2-M644G msh2Δ* cells.

### Pol ε proofreading domain mutations increase the frequency of insertions

Analysis of the number of insertions/deletions accumulated in the absence of MMR, which would otherwise efficiently repair them, revealed that Pol ε exo^−^ introduces insertions ~4.5 times more frequently than wild-type Pol ε does, and that this is further increased two-fold in the presence of Pol ε–L439V (**Fig. 4A**). Analysis of the type of base inserted revealed that in both cases, mutant Pol ε is mainly responsible for the introduction of +T and +A, which overall represent 70-80% of all insertions (**Fig. 4B,** as opposed to ~31% for *msh2Δ* strains). Since these insertions likely arise from the role of Pol ε as the leading strand replicase[23], replication-strand bias analysis suggests that Pol ε – L439V is mostly responsible for +A insertions (**Figure 4C**). Moreover, these +A insertions on the leading strand tend to occur in the context of a 3-6 nucleotide T homopolymer in the template strand (**Fig. 4D**), suggesting that they originate from polymerase slippage.

**Figure 4.**
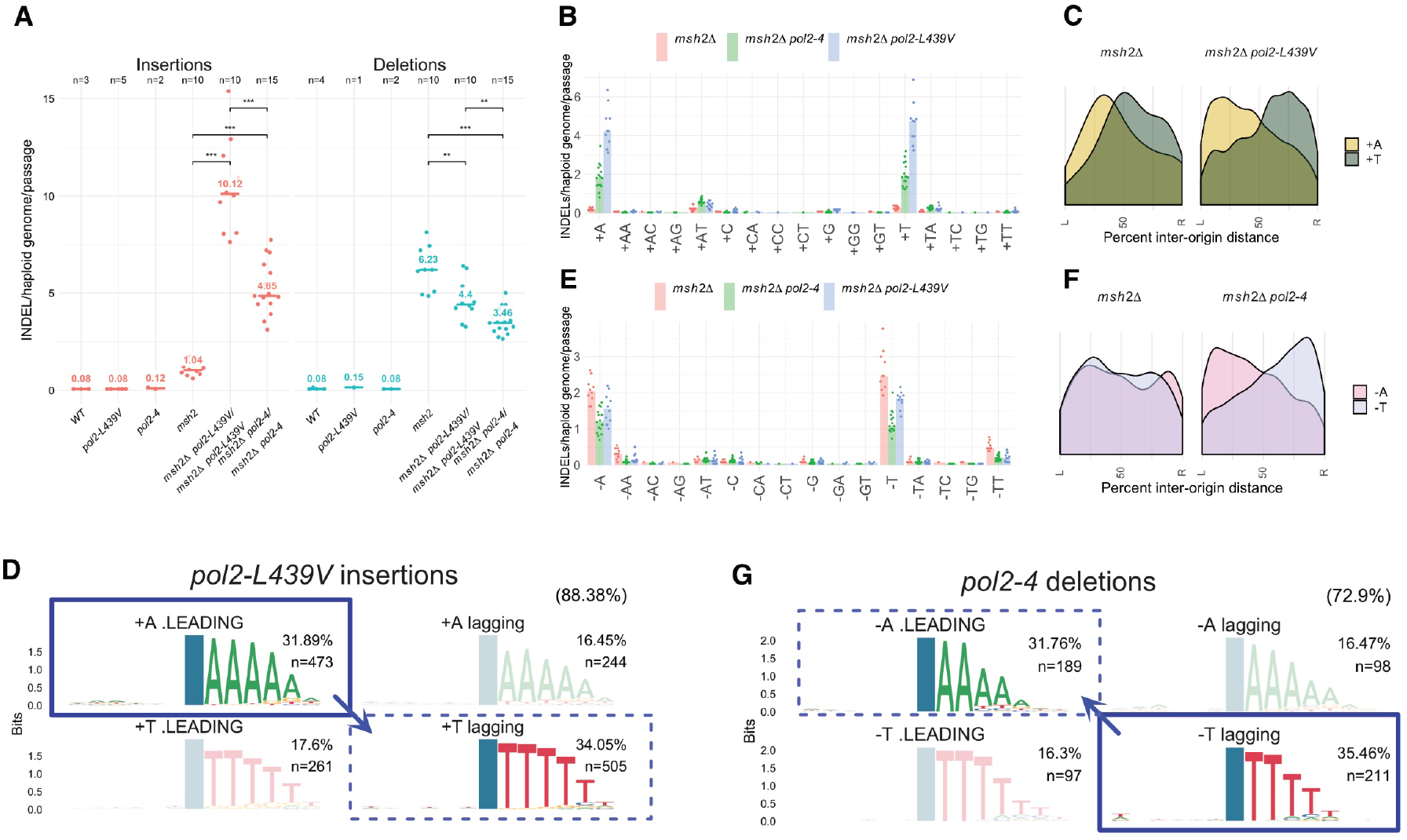
PolE ultra-mutators frequently introduce +A insertions opposite T homopolymers. **A.** Rate of insertions and deletions introduced by Polε *exo*^−^*(pol2-4)* and the ultra-mutator Polε L439V *(pol2-L439V)* in the absence of functional mismatch repair *(msh2Δ).* B. Rate of insertion of different mono-or di­ nucleotides. **C.** Geographical distribution of the density of the two most common types of insertion in relation to replication origins. **D** Sequence context in which +A/+**T** mutations occur in relation to the replication strand in which it occurred; the numbers in brackets indicate the prevalence of this class compared to all insertions. The conserved homo-polymer context always appears on the 3’ side of the mutation because INDELs were left-aligned during variant normalization. Solid boxes indicate that the insertions observed on the leading strand are introduced directly or indirectly by Pol ε. Dashed boxes indicate that the complementary sequence context is observed in lagging strand regions. E. Rate of deletion of different mono-or di-nucleotides. F. Geographical distribution of the density of the two most common types of deletion in relation to replication origins. **G.** Sequence context in which -A/-T deletions occur in relation to the replication strand in which it occurred; the numbers in brackets indicate the prevalence of this class compared to all deletions. The conserved homo-polymer context always appears on the 3’ side of the mutation because INDELs were left-aligned during variant normalization. Solid boxes indicate that the deletions observed in *pol2-4* strains are likely introduced on the lagging strand, possibly by Polδ. Dashed boxes indicate that the complementary sequence context is observed on the leading strand.

### Pol ε proofreading activity appears to be mutagenic

Differently from what we have observed for insertions, MA analysis revealed that the production of short, mostly single-base, deletions was ~50% lower in *pol2-4* cells compared to wild-type *POL2*. This suggests that Pol ε exonuclease activity is directly responsible for essentially half of the deletions produced during normal DNA replication in the absence of MMR which would otherwise repair them. The Pol ε – L439V mutant showed an intermediate deletion rate phenotype, possibly because this mutant could retain partial exonuclease activity, as it was shown for Pol ε – P301R. Analysis of the spectrum of deletions implied that a wild-type replisome largely introduces -T, -A, -TT, and -AA deletions, and that Pol ε exonuclease activity contributes to roughly half of these (**Fig. 4E**). Analysis of replication-strand bias also revealed that the frequent -A and -T deletions do not normally show any discernible strand preference. Inactivation of Pol ε exonuclease activity, however, led to a strong bias for -A deletions on the leading strand and corresponding -T deletions on the lagging strand (**Fig. 4F**). These results suggest that both Pol ε exonuclease activity and an unidentified process on the lagging strand (possibly Pol δ exonuclease activity) produce frequent -T deletions, giving rise to no overall replication-strand bias. In *pol2-4* cells, however, it appears that the leading strand branch of this mutagenic pathway is inactivated, generating a -A deletion bias that is the reflection of the -T deletions produced on the lagging strand (**Fig. 4G**).

## DISCUSSION

DNA-replication associated hypermutation is a known driver of colorectal and endometrial cancers, whether arising from mismatch repair (MMR) inactivation or from DNA polymerase ε or δ exonuclease domain mutator (EDM) alleles. While it is clear how MMR inactivation increases mutagenesis, establishing the source of mutations generated by EDM alleles has proven more difficult. The yeast benchmark for EDM alleles, *pol2-P301R*, generates mutations at a much higher rate than the corresponding exonuclease-dead strain (*ultra*-mutator phenotype) [13], while retaining part of the wild-type exonuclease activity [15]. Our results now show that the *ultra*-mutator phenotype is shared by some other yeast Pol ε EDM alleles orthologous to cancer-associated Pol ε mutations, and that increased mutation accumulation in *pol2-ultra* cells compared to *pol2-4* cells also occurs in the absence of MMR activity, strongly suggesting that differential repair of mismatches is not a major source of hypermutation.

Studies of Pol ε protein structure have revealed that the P301R mutation creates a positively charged surface that likely hinders a 3’ mismatched primer end from properly accessing the proofreading domain and could, thus, favour its extension [15,16]. Strikingly, the second strongest mutator that we identified (*pol2-M459K*) also introduces a positive charge in the same area (**Additional File 1: fig. S4**), while the weaker *ultra*-mutator alleles could also hinder proper DNA strand placement, perhaps by altering Pol ε structure more subtly.

The ultra-mutator phenotype could be explained if mis-pairs observed in cells lacking Pol ε exonuclease activity are not caused by impaired removal of a mis-incorporated base, but rather mainly arise from an active mutagenic process driven by Pol ε and whose intensity is heightened by Pol ε–*ultra* mutants. A prediction of this model is that Pol ε *ultra* and Pol ε exo^−^ would produce the same pattern of mutations; and, indeed, the mutational profiles, strand bias, and sequence contexts in which mutations occur in *pol2-ultra* and *pol2-4* cells alleles are virtually indistinguishable from each other. In this scenario, Pol ε could contain a “mutagenic proofreading” activity, normally suppressed by the presence of exonuclease activity, and activated by *ultra*-mutator alleles. We suggest that this activity arises from the backtracking of Pol ε in the first stage of proofreading. In this model, under normal conditions, this movement melts the primer-template junction until the nascent 3’ end has been inserted in the exonuclease domain for hydrolysis (**Fig. 5A**). In the absence of hydrolysis — or even more so when access to the exonuclease active site has been blocked by *ultra* mutations — the nascent strand would prevent this movement. In these situations, backtracking could still occur if the nascent strand were to shift backwards and extrude a base further upstream (**Fig. 5B**), an activity that would result in the generation of insertions, especially after A:T homopolymers that are easier to melt than G:C ones because of their weaker bonding. At this point, further polymerisation would require the mis-inserted base to form a proper pair with the T template, and thus would occur only when an adenine was mis-inserted in the first place (**Fig. 5C**). In agreement with this model, we found that in the absence of MMR, *pol2-4* cells and to a greater extent *pol2-L439V* cells, frequently accumulated +A insertions when replicating through T-homopolymers. Furthermore, when two adenines follow the T-homopolymer template, the first one becomes a hotspot for A:A mis-pairs. In our model, this would arise from the newly synthesised strand sliding forward after the first base post-mismatch has been introduced. This would restore full base pairing, converting the +A insertion into a A:A mispair and generating the observed T>A transversions (**Fig. 5D**).

**Figure 5.**
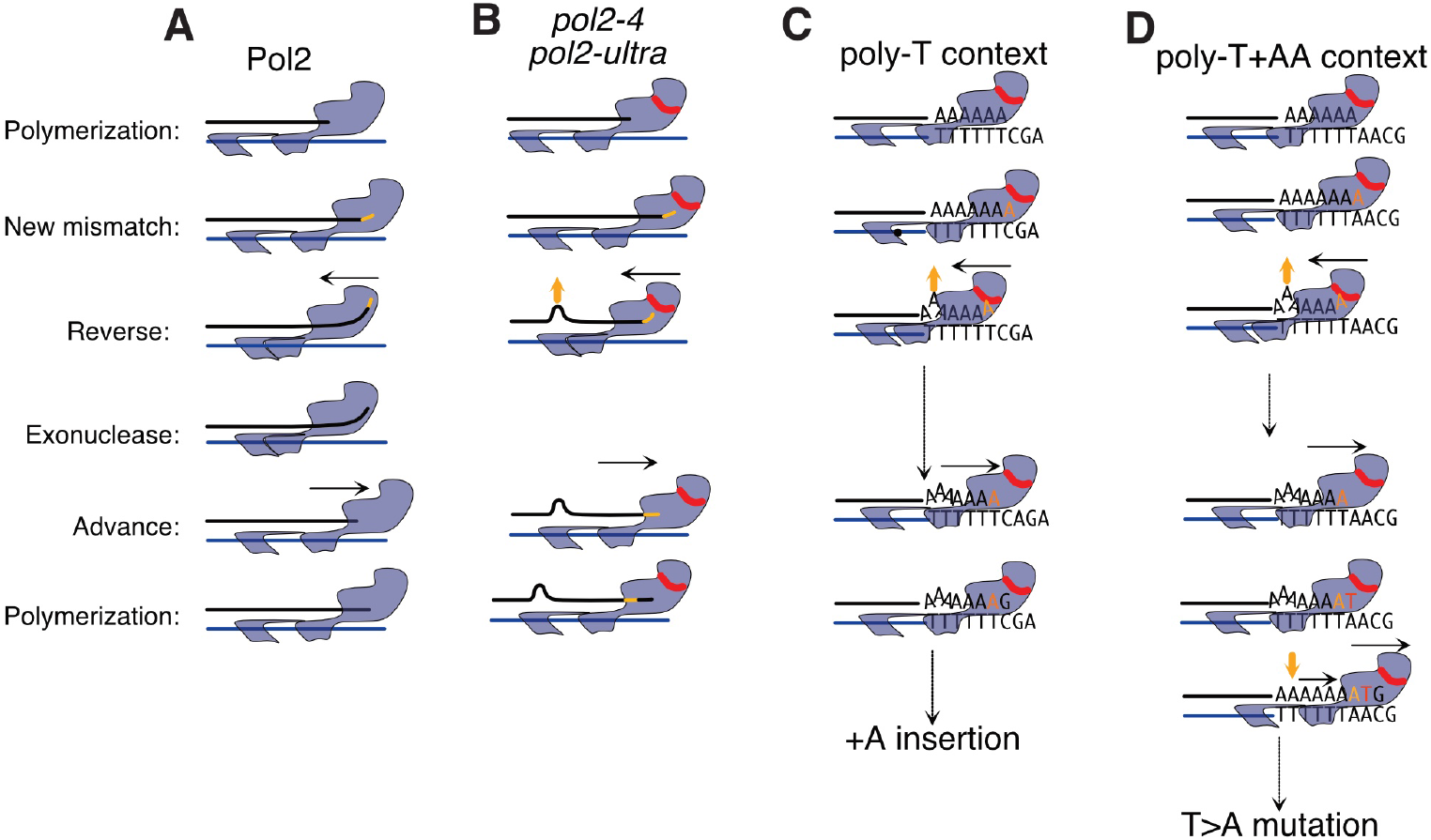
Model for the origin of mutations specifically introduced by Pol ε exonuclease domain mutants. **A.** In the presence of a terminal mismatch (yellow segment), Pol ε backtracks (black arrow) and repositions the terminal-mismatched primer in the proofreading domain; after hydrolysis the primer is re-annealed and extended. **B.** In the presence of exonuclease-inactive mutants or *ultra-mutator* mutants backtracking is blocked unless the newly synthesised strand melts, shifts back and extrudes one base (yellow arrow). **C.** Melting and extrusion can only occur on homopolymer templates (TTTTT), where the shift does not alter correct base-base pairing between template and nascent strand. If the mis-inserted base is an adenine (yellow A), this can pair with the last T of the template and reconstitute a proper base pair that can be extended, generating insertions. **D.** If a T homopolymer is followed by two A’s, then a T would be inserted in front of the first A (red T), but would then move in front of the second, when the base extruded in the homopolymer is re-annealed.

Given the sequence and mis-insertion requirements needed, the above-described events should be comparatively rare; and indeed, insertions represented only ~10% of all the mutations we observed, while T>A transversions accounted for less than 2%. With the exception of these classes of mutations, we found that the sequence context of the remaining mutagenic channels closely resembled the context of mutations deposited by a low-fidelity Pol ε mutant carrying a mutation in the polymerase domain (*pol2-M644G*). The similarity between mutations introduced by Pol ε exonuclease and polymerase domain mutants strongly suggests that they foster the same mechanism of mutagenesis. While this model does not exclude that another polymerase could be responsible for introducing mis-pairs on the leading strand after Pol ε stalling (for example through Pol α–mediated re-priming), it does suggest that the majority of mutations introduced in *ultra*-mutator cells are directly introduced by Pol ε. In this regard, the mutagenic pattern shared by polymerase and exonuclease domain mutants could arise from the increased dNTP levels observed in *pol2-M644G* cells, which has been shown to contribute to its mutator phenotype [29]. This could also explain why Pol ε – P301R does not show the same heightened level of mutagenesis *in vitro* as compared to Pol ε exo^−^.

In conclusion, our findings have provided further insights into how cells normally guard against mutagenesis during DNA replication, and how specific point mutations in replicative polymerases affect their function to heighten mutation rates and lead to distinctive mutational signatures. Given that DNA replication and replicative polymerases are highly conserved throughout evolution, the effects and mechanisms that we have described likely also operate in cancers with orthologous polymerase mutations.

## MATERIALS AND METHODS

### Yeast strains and plasmids

For the mutations of interest, identified from the literature as amino acid changes, sequence alignment with Clustal Omega version 1.2.1 was carried out to determine the orthoogous *S. cerevisiae* residues. Uniprot sequences used for alignment were *Homo sapiens* POLE (Q07864) and *S. cerevisiae* POL2 (P21951). All *S. cerevisiae* strains used were derived from the laboratory strain W303 (*leu2-3,112 trp1-1 can1-100 ura3-1 ade2-1 his3-11,15 RAD5*). Polymerase mutations were created by cloning an N-terminal *POL2* PCR fragment into pRS306 and generating the mutations of interest by site directed mutagenesis using the QuickChange Lightning Kit following manufacturer’s instructions (Agilent Technologies). Polymerase mutants were introduced into *MAT***a** haploid *S. cerevisiae* W303 strains, resulting in a full-length copy carrying the mutation and a non-mutated, truncated N-terminal fragment. Haploid *pol2* mutants were then mated to a wild-type isogenic *MATalpha* strain to generate heterozygous diploid mutant strains. Deletion of *MSH2* was introduced in wild-type W303 by one-step gene disruption. Disruptions were confirmed by PCR and whole-genome sequencing. Haploid double mutants *pol2 msh2Δ* were recovered by mating, sporulation, tetrad dissection and analysis. The genotypes of strains are described in Supplementary Table 1.

### Growth rate and mutation assays

The growth rates of heterozygous diploid polymerase mutant strains were assessed by growing cultures to stationary phase, diluting them into rich medium and growing for 450min. Growth was assayed by measuring absorbance at 595nm wavelength. To determine mutation rates at the *LYP1* locus, single colonies were excised from agar plates, inoculated in rich medium, and grown to saturation. Cell cultures were subsequently diluted 1:100,000 and plated on YPAD (1% yeast extract, 2% peptone, 2% glucose, 40 mg/l adenine) plates, or plated (50 μl – 250 μl) without dilution on SD (Synthetic Defined media) -LEU +Thialysine (50 mg/l) plates. Different amounts were used for different strains to obtain countable plates. Mutation rates were calculated as described in [30].

### Single-cell bottleneck propagation

Heterozygous diploid strains were grown on solid media at 30°C and propagated for 26 passages, every 2-3 days, through single-cell bottlenecks by means of repetitive isolation and single-colony picking. Cells were estimated to have undergone ~20 generations per passage to generate a colony of 10^6^ cells (approximately 500 generations during the entire propagation). Colonies were picked randomly to avoid bias towards adaptive or deleterious mutations (with the exception of lethal mutations). In each experiment, each strain was propagated in 18 parallel lines. Deviations from this number due to failures to sequence or to confirm the presence of the mutation at the end of the propagation are denoted in the relevant figures. Haploid strains were propagated for 13 passages. Standard YPAD non-selective rich medium was used. In each experiment, whole-genome sequencing of two random colonies for each strain was attempted at passage 0 and at the end of the propagation. Only mutations observed in both colonies (where a second colony was available), and absent from passage 0, were retained for further analysis.

### Small-population bottleneck propagation

Haploid and heterozygous diploid polymerase mutant strains were propagated in a 1536 plate format in a non-selective complete synthetic medium (0.14% YNB, 0.5% ammonium sulphate, 0.077% complete supplement mixture [ForMedium], 2% (w/v) glucose and pH buffered to pH 5.8 with 1% (w/v) succinic acid). Plates were replicated using a ROTOR Robot (Singer Ltd, UK) and 1536 short pin pads every 2-3 days for 40 passages, through bottlenecks estimated to contain ~10^4^ cells. Effective population size is, however, likely to be smaller due to the population structure of cells in a colony. The number of cells in the bottleneck was calculated by estimates of pixel intensities using light transmission and conversion of pixel intensities into cell counts by calibration to a flow cytometry-based reference[31]. In these conditions, wild-type BY strains undergo ~6 doublings per growth cycle (passage) suggesting that cells underwent ~240 generations over the duration of the experiment. Final populations were streaked for single colonies and the whole genome 18–26 isolates per strain was sequenced. After reassigning strains to the correct genotypes and ploidy 12–38 isolates per strain were analysed.

### DNA extraction, library preparation and whole genome sequencing

Genomic DNA extractions and library preparations were carried out as previously described [32]. Libraries were sequenced using either HiSeq 2000 or HiSeq X (Illumina) to generate 125bp or 150 bp paired-end reads, respectively.

### Reference genome alignment

Sequencing reads were aligned to the *S. cerevisiae* S288c (R64-1-1) reference genome using BWA mem (-t 16 -p -T 0) and duplicates were marked with bamstreamingmarkduplicates (biobambam2 2.0.50) and stored in CRAM format (primary data). From these, reads were extracted with *samtools fastq* and subsequently re-aligned to a modified reference genome in which repetitive DNA regions were hard-masked and moved, as single-copy sequences, to *ad hoc* artificial chromosomes. Duplicates were marked with *bamsormadup SO=coordinate fixmate=1*.

### Confirmation of strain genotypes

Samples were automatically checked for their expected polymerase genotype using the script *deletion_check.pl.* Briefly, for point mutations the DNA sequence from the triplet coding for the residue in question was extracted from the sequencing data, translated and compared with the expected. Deletions and genetic mating type were determined as previously described [32]. Ploidy was determined a posteriori, based on the distribution of the observed allelic frequencies (AF). Strains displaying a majority of alleles with AF ~0.5 were classified as diploid, while strains in which the majority of alleles had an AF of ~1 were classified as haploid.

### Variant calling, consequence annotation and filtering

SNVs and small insertions/deletions (INDELs) were identified chromosome by chromosome using *samtools mpileup* (v.1.9), with the following options: *-g -t DP,DV -C0 -p -m3 -F0.2 -d10000*, followed by *bcftools call -vm -f GQ* (v.1.9). All mutations from each chromosome were merged with *bcftools concat*. All variants were annotated with the Ensembl Variant Effect Predictor (VEP; v95.3). INDELs were subsequently normalised with *bcftools norm -m-both --check-ref e* and sorted with *bcftools sort*. Low quality variants were flagged with *bcftools filter* with the following options *-m + -e ‘INFO/DP<10’ -e ‘FORMAT/DV<3’ -e ‘TYPE=\“snp\” & QUAL<100’ -e ‘TYPE=\“indel\” & QUAL<30’ -e ‘FORMAT/GQ<40’ -g 7*. Variants present in control samples were subsequently removed with *bcftools isec -w1 -C {sample_file} {control_file}*.

### Further mutation filtering

SNV mutations were further filtered on the QUAL value and their prevalence across different sequencing samples. Given the relatively low number of mutated positions compared to the genome size, the vast majority of mutations are expected to be unique in different MA lines in single-cell bottleneck experiments, and shared mutations are likely to originate from systematic sequencing errors. Taking this into account, we removed mutations whose quality was below an arbitrary threshold that grows linearly with the prevalence of the mutation in different samples (**Additional File 1: fig. S5**), thus excluding mutations that are frequently observed and of lower quality. In small-population bottleneck experiments, many mutations are shared between different colonies from the final population, because of their shared ancestry. For this reason, a similar, less stringent threshold was used. Filters were designed to remove approximately 1-10% of all mutations. A similar rationale was used to filter INDELS. Small changes in the filtering parameters do not substantially alter the results of the subsequent analyses.

### Analysis of mutation numbers

The total number of SNPs/INDELs for each sequencing sample was calculated by counting the number of mutations passing all filters. In single-cell bottleneck propagations two colonies per MA line were sequenced and only mutations observed in both colonies (and absent from passage 0) were retained for further analysis; where a second colony was not available because of sequencing failure, all mutations in the only available colony were retained. For small population bottleneck propagation all mutations present in each colony from the final population were considered. Mutation rates are given in terms of SNV(INDEL)/haploid genome/passage and converted to SNV/generation/bp assuming a haploid genome size of 12,071,326 bp and 20 generations from single cell to colony.

### Analysis of mutation types

In small-population propagation experiments, one single mutagenic event is likely observed in more than one colony picked from the final-population. Thus, mutations derived from small-population experiments were initially grouped by MA line and, in each line, when the same mutation was observed in more than one colony, only one instance was retained. Mutations from single-cell and small-propagation experiments were then pooled for the purpose of the subsequent analyses. Analysis of the frequency of different SNV classes was carried out by grouping the mutations by genotype (irrespectively of the type of propagation or ploidy), counting the number relevant mutations, and summing complementary pairs (e.g. A>C + T>G).

### Analysis of replication strand bias

The relative position of each mutation with respect to the nearest replication origin was calculated in two steps. First, a replication model was built using the coordinates of replication origins obtained from OriDB (http://cerevisiae.oridb.org; only using “confirmed” origins); location of each origin was calculated as the midpoint of the ARS region; termination points were arbitrarily defined as the midpoint of each inter-origin span; leading-strand regions were defined as regions comprised between an origin and the termination point to its immediate right; lagging-strand regions were defined as regions comprised between an origin and the termination point to its immediate left. Second, each mutation was localised to an inter-origin span (thus, mutations located before the first origin or after the last origin of each chromosome were discarded); the distance between each mutation and the origin to its immediate left was calculated, and normalised for the size of the inter-origin span in which the mutation was located, so that a distance of 50% coincides to the midpoint termination zone, and a distance of 100% coincides to the subsequent origin. To avoid over-weighting shared mutations originating from small-population bottleneck experiments, only a distinct set of mutations was considered. The density of each mutation type was then plotted as a function of the relative distance of mutations from the origin to the immediate left.

### Analysis of mutation patterns and comparison with mutation signatures

Mutational patterns were obtained by calculating the frequency of the 96 trinucleotide contexts (channels) in which mutations belonging to one of the six main classes (C>A, C>G, C>T, T>A, T>C, T>G) occurred. The remaining mutations (G>T, G>C, G>A, A>T, A>G, A>C) were reverse complemented along with their context and assigned to the appropriate channel. The frequency of each channel was then normalized by the relative abundance of each trinucleotide in the yeast genome.

Comparison with mutational signatures identified in cancers was calculated using cosine similarity [33] and the *cos_sim_matrix* function of the *MutationalPatterns* R package [34]. Cancer signatures were obtained from https://cancer.sanger.ac.uk/cancergenome/assets/signatures_probabilities.txt.

### Analysis of mutation sequence context

After extracting the context (5 nucleotides) in which each mutation occur, mutations were classified as leading or lagging strand mutations depending on where they occurred in relation to origins of replication and presumed termination points (see Analysis of replication strand bias). To exclude as much as possible mutations introduced by Pol ε synthesising DNA beyond the midpoint of each replicon, only mutations occurring in the first and last third of each replicon where considered. Sequence logo images from the context of each mutation class were obtained with the *ggseqlogo* R package[35].

## Acknowledgements

We thank D. Robles-Espinoza, M. del Castillo, N. Geisler, M. Rashid, K. Wong, I. Martincorena, L. Alexandrov, V. Iyer, C. Bradshaw, T. Keane, and members of the SPJ laboratory for advice and discussion. This research was supported by: Wellcome Strategic Award 101126/Z/13/Z (COMSIG); Wellcome Investigator Award 206388/Z/17/Z; Wellcome PhD Fellowship 098051 to MH; Cancer Research UK Programme Grant C6/A18796; Cancer Research UK C6946/A24843 and G2244/A21717, Swedish Research Council grants 2014-6547 and 2014-4605, and Wellcome WT203144 Institute Core Funding.

## Data Availability

Primary sequencing data has been deposited at the European Nucleotide archive (https://www.ebi.ac.uk/ena/browser/home) with the accession IDs indicated in Supplementary Table 2.

## Author contributions

The project was conceived by SPJ, DA, IS, FP, and JW. Manual and automatic propagations were carried out by MH and EA-P respectively. DNA extractions were carried out by MH. Data analysis was carried out by FP and MH. The manuscript was written by FP and SPJ, with contributions from JW, MH, DA and IS.

---

**Supplementary Figure 1.**
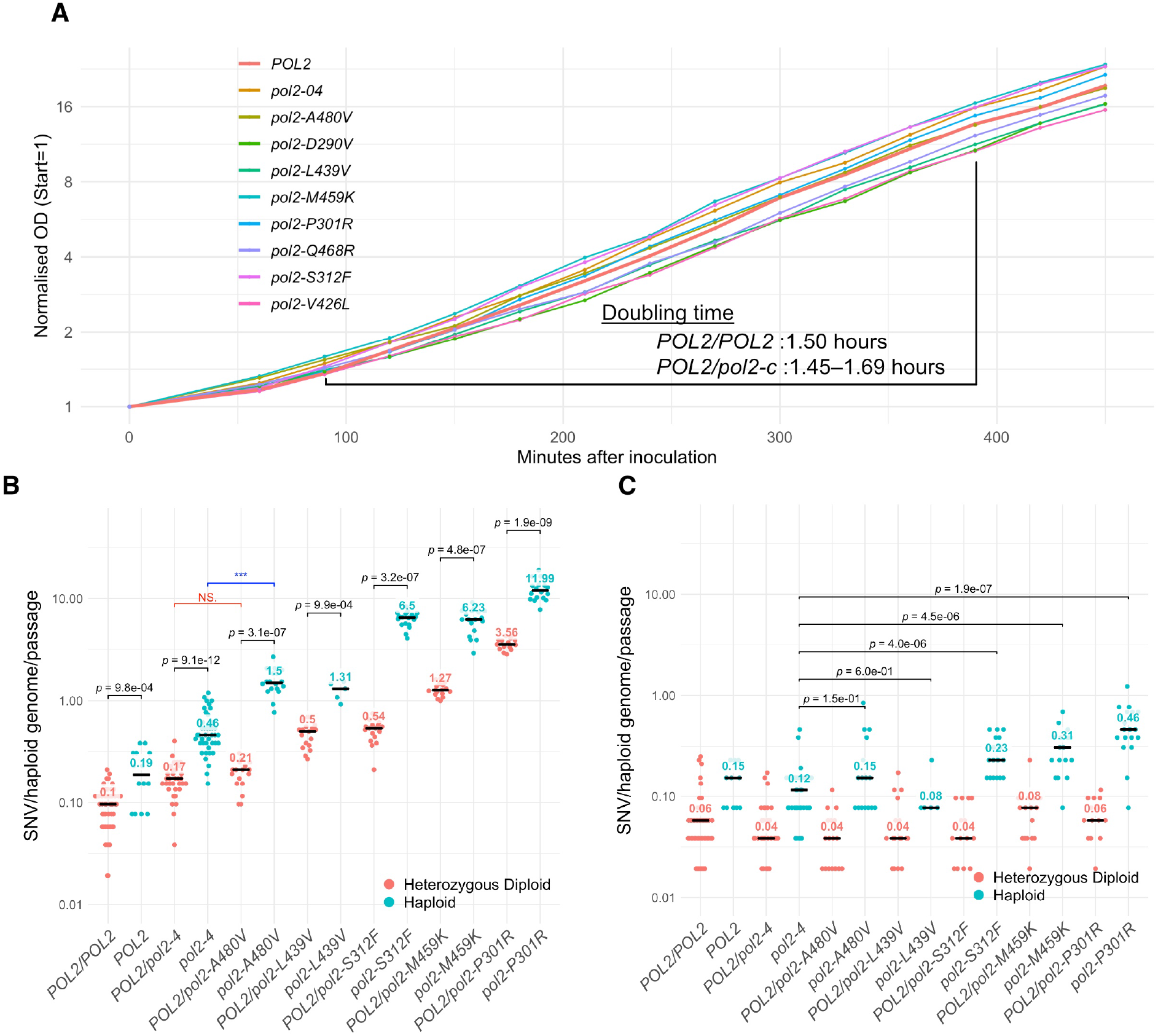
Comparison of mutation rates in haploid and heterozygous diploid Pol ε mutants. **A.** Growth curves of different heterozygous diploid polymerase mutants. At the time points in which OD exceeded 0.8 the OD was measured from serial dilutions and the theoretical OD calculated multiplying the measured OD by the dilution factor. All values were then scaled so that the OD at the initial time point = 1. **B, C.** Rates of SNV and INDEL accumulation in strains carrying the indicated *POL2* alleles in haploid and heterozygous diploid cells propagated through single-cell bottlenecks for 26 (diploid) or 13 (haploid) passages (~500 or ~250 generations). Each independent mutation accumulation line is indicated by a dot. Statistical test: Mann-Whitney-Wilcoxon.

**Supplementary Figure 2.**
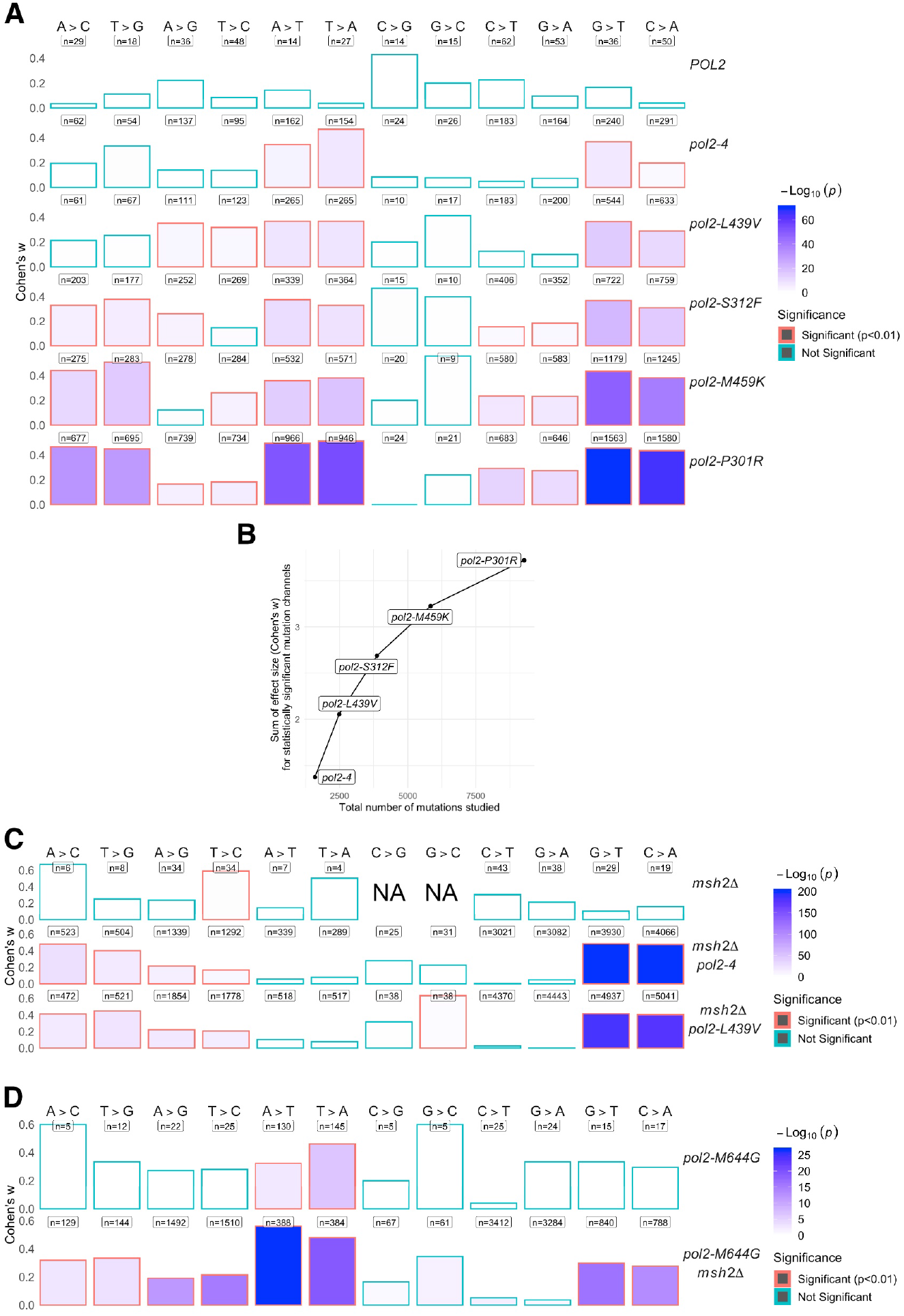
χ^2^ test for deviation from uniformity of the distribution of the inter-origin position of mutations. **A.** Graphical representation of how significantly the distribution of each mutation type on the leading and lagging strand deviates from a uniform distribution. For each mutation type in each genotype, mutations were binned into leading or lagging strand based on their relative inter-origin position. P-values were obtained from χ^2^ tests and corrected for multiple testing using the Holm­ Bonferroni method *(n* = 12). The height of each bar is proportional to the effect size of the deviation from uniformity (Cohen’s *w*); the total number of observations is indicated above each plot; the filling color is proportional to −log_10_(p) and mutation types significantly (p < 0.01) deviating from uniformity are outlined in red. Primary data from Fig. IF. **B.** Correlation between the sum of the effect size of deviation from uniformity for each mutation type in each genotype and the total number of mutations studied by genotype. **C.** Same as panel A; Primary data from Fig. 2D. **D.** Same as panel A; Primary data from Fig. 2E.

**Supplementary Figure 3.**
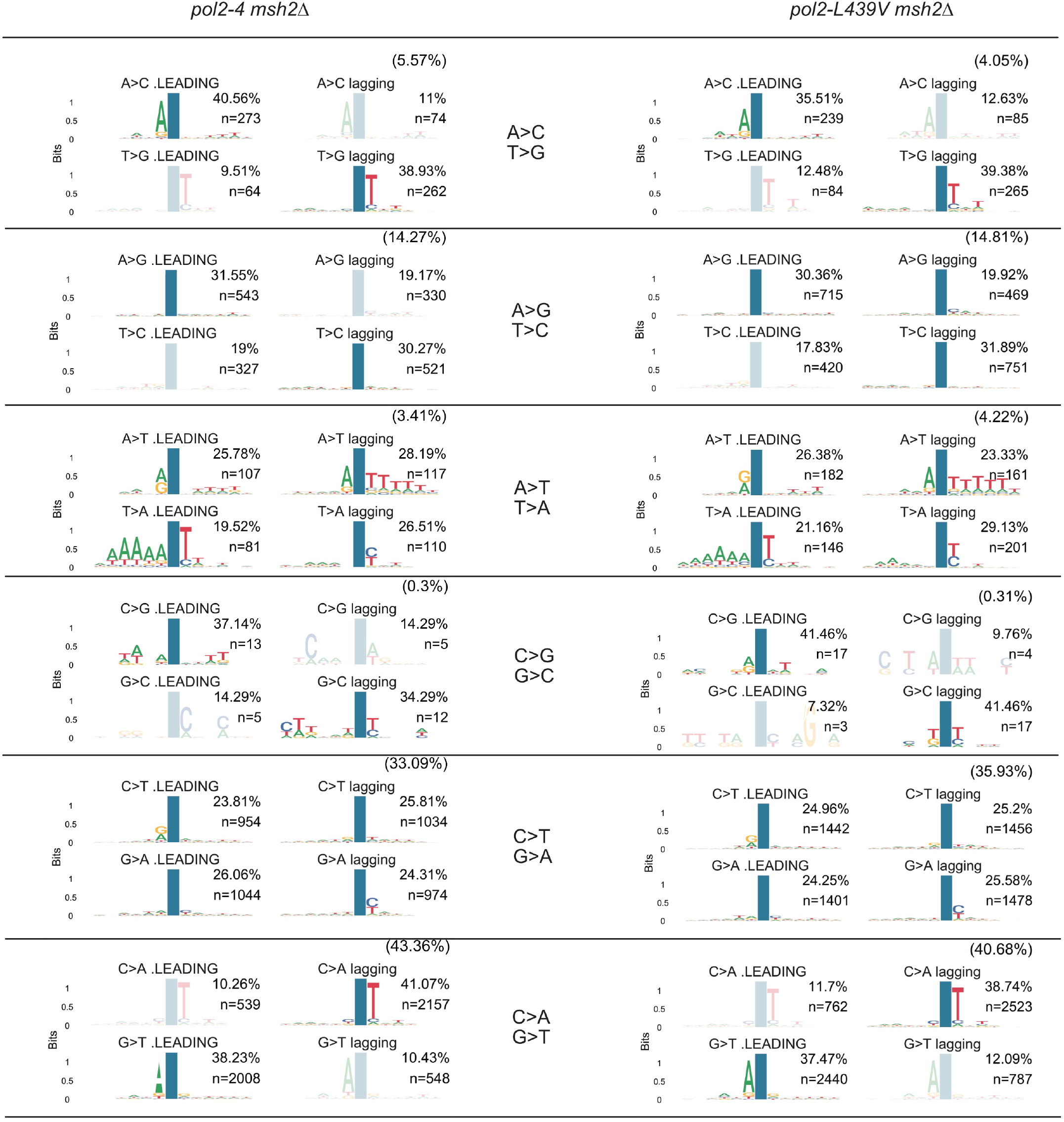
Sequence context of the mutations generated in cells with Pol ε exo^−^ or Pol ε ultra mutants. Sequence context in which each mutation class occurs in cells carrying Pols exo^−^ *(pol2-4)* or Pols ultra-mutator *(pol2-L439V),* in the absence of functional MMR *(msh2Δ)* Mutations were categorised by type and by wether they occurred in regions of the Watson strand synthesised as leading or lagging strand. The frequency of each mutation class is indicated in brackets. Whithin each class the frequency of the four types is indicated in each panel.The position of the mutation is indicated by a blue column. Transparent classes indicate that these mutations are likely artefacts created by leading strand replication not precisely terminating at the inter-origin midpoint and/or by some origins being passively replicated.

**Supplementary Figure 4.**
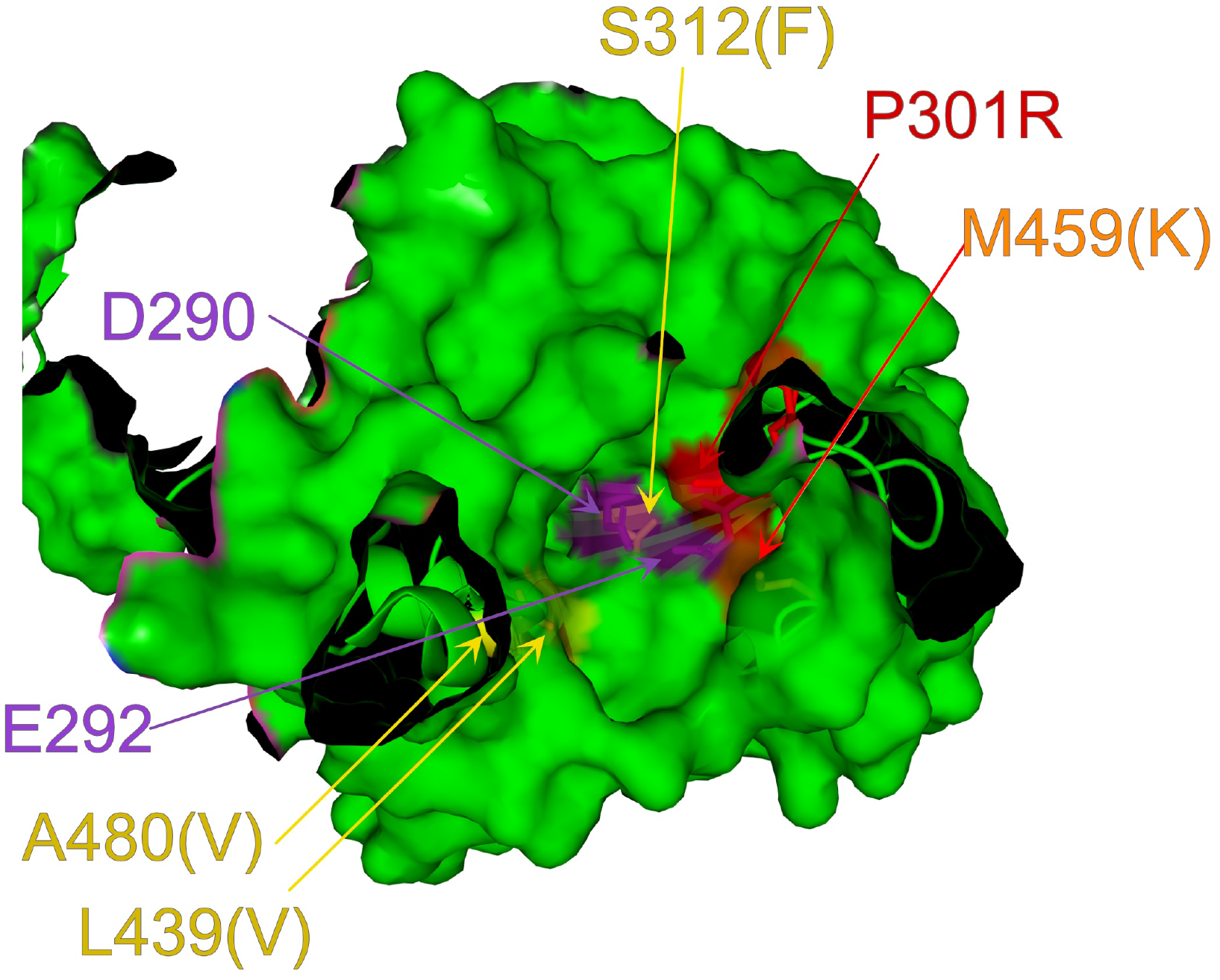
Location of the residues mutated in Pol ε proofreading domain. Structure of the proofreading domain (residues 271-491) of DNA polymerase ε with the P301R mutation (6G0A). The residues that when mutated confer an *ultra*-mutator phenotype have been highlighted and the mutation is indicated in parentheses. The second strongest mutator M459K would introduces a positive charge next to the positive charge introduced by the strongest mutator P301R. S312F, L439V and A480Vare weak *ultra*-mutators.

**Supplementary Figure 5.**
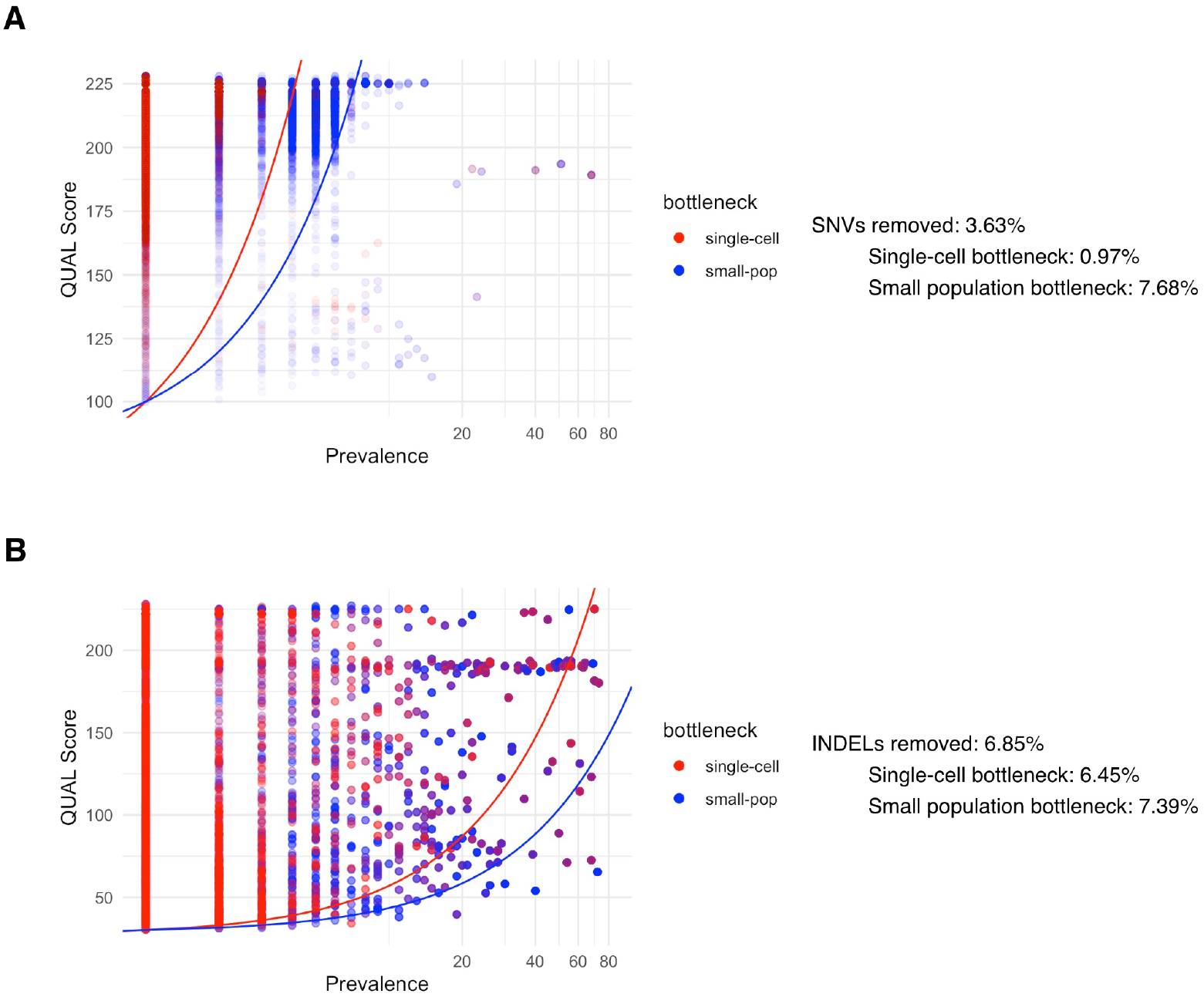
Additional quality filtering of mutations. Single nucleotide variants **(A)** and insertion-deletions **(B)** are plotted as a function of the number of times they have been observed in the dataset (Prevalence) and their overall quality score (QUAL). The filtering thresholds used for single-cell and small-population bottleneck experiments are marked by a red and blue curve, respectively. The number of mutations discarded for each class is indicated next to each plot.

